# The Georgetown Reading in Aging Neuroimaging Dataset (GRAND): Reading and multimodal MRI data in older adults

**DOI:** 10.64898/2026.05.18.725986

**Authors:** Elizabeth J. Anderson, Ryan Staples, Sara M. Dyslin, Elizabeth H.T. Chang, Alycia B. Laks, J. Vivian Dickens, Devna Mathur, Sachi Paul, Elizabeth Dvorak, Peter E. Turkeltaub

## Abstract

Reading is a critical skill in modern society. Most research on reading is conducted in school age children or young adults. However, acquired brain disorders often affect reading ability, and these disorders tend to occur in older adults. It is therefore critical to examine the normative distribution of reading behavior and the brain basis of reading in older adults. Here, we provide trial-wise single word and pseudoword oral reading and lexical decision data, as well as structural, functional, and diffusion-weighted MRI data from 116 neurotypical adults aged 22 to 84 years (mean = 59). Accuracy, response times, and errors are provided for corpora that are parametrically modulated in frequency, imageability, and regularity for real words and consistency of spelling-sound mapping for pseudowords. This dataset includes both minimally processed behavior (trial-wise data) and MRI data, and participant- and item-wise summary metrics and processed MRI data. These data serve both as a normative sample for reading behavior in older adults, but also as a valuable resource for identifying novel brain-behavioral relationships.

## Background & Summary

Reading is a vital skill in modern societies. Reading impairment is thus a serious disability, leading to negative impacts on life activity, social participation, and quality of life^1–3^. Accurate assessment of reading disorders is needed clinically for assignment to treatments and scientifically for identification of the cognitive processes that are affected by neural damage. Critically, assessment of reading impairments depends on an age-matched reference sample: aging is associated with a general psychomotor slowing that is apparent in both oral reading and lexical decision^4,5^. Some evidence also suggests that older adults are more sensitive to lexical variables, such as word frequency, and less sensitive to sublexical variables as compared to younger adults^6,7^.

This dataset was collected as a normative reference for studies of stroke-induced language and reading impairment. Diagnosis of alexia (acquired reading impairment) is typically accomplished by assessing performance on single-word language tasks. We thus assess reading ability using two tasks, oral reading and lexical decision, to examine visual word processing with and without production.

The stimuli are single-syllable words and pseudowords, manipulated across several psycholinguistic dimensions. We provide 1) minimally processed trial-level accuracy, response time, and (for oral reading) error codes and transcriptions, 2) participant-wise averages across all metrics, and 3) item-wise averages across all metrics. We have previously used subsets of this data as age-matched normative reference samples for studies examining the role of phonological^8^ and semantic impairment^9^ in oral reading, and for assessing the overlap and divergence of neural processes supporting oral reading and lexical decision^10^.

We also provide high resolution T1-weighted and fluid-attenuated inversion recovery (FLAIR) MRIs, high angular resolution diffusion weighted imaging (HARDI), and functional MRI (fMRI) data from a written semantic relatedness decision task designed to robustly assess language processing across a range of ability levels^11^. We provide fMRI data in both a minimally preprocessed time-series format, for researchers who intend to use their own processing pipeline, and the participant-level critical contrasts that are the output of our processing pipeline. We also provide whole-brain structural connectivity matrices derived from the HARDI data.

These data constitute a normative assessment of single word reading and of the language network in typical older adults. As such, the data provide an opportunity to assess the cognitive effects of disease and injury processes common in older adults. Further, the data are appropriate for questions examining how reading ability is related to language activation in the brain. The data are amenable to activation and task-related connectivity analyses. Beyond the domains of reading and language, the T1-weighed, FLAIR, and HARDI data can be used to address questions related to brain structure and connectivity during aging more broadly.

## Methods

116 participants (Table 1) completed four reading tasks as part of larger batteries for one of two study protocols, labeled BUILD (Brain-based Understanding of Individual Language Differences after stroke, ClinicalTrials.gov ID: NCT04991519, PI: Turkeltaub, N = 69) or ReadMap (Reading in Stroke Alexia and Typical Aging, ClinicalTrials.gov ID: NCT06700005, PI: Turkeltaub, N = 47). Both studies were approved by the Georgetown University Institutional Review Board, and informed consent was obtained prior to participant testing. Participants were 18 years or older, had no major neurological or psychiatric disorders, learned English before the age of eight, and scored above the Montreal Cognitive Assessment (MoCA)^12^ total score cutoff adjusted for age, ethnicity, and race^13^.

**Table 1.** Participant Demographics.

|  |  |
| --- | --- |
| Age in years | 59.52 (sd: 12.74; range: 22.42 – 84.11) |
| Sex (Male/Female) | 56/60 |
| Race (Asian/Black/White/Multiple) | 2/40/71/3 |
| Education (Years by Degree) | 16.99 (sd: 2.31; range: 12 – 21) |

*Note.* Values are mean (standard deviation).

Task structure and stimuli were largely similar across protocols, with any differences noted below. Items were displayed in the center of a Dell Inspiron 17′′ 2-in-1 touchscreen or a Lenovo Yoga 7i 16” 2-in-1 touchscreen in the tented position for a maximum of 10 seconds. Each trial began with a fixation cross, followed by a 300ms 460Hz tone for the oral reading tasks, and then the presentation of the item. Participants were prompted to touch the screen to continue after 10 seconds if they had not advanced to the next item. For the oral reading tasks, RTs were calculated from the offset of the tone to the onset of speech.

### Real Word Oral Reading

Participants read aloud 200 monosyllabic English words manipulated systematically for frequency (low vs. high), spelling-to-sound regularity (regular vs. irregular), and imageability (low vs. high) as previously described in Staples et al., 2025^9^. The corpus thus included eight different word types (e.g., high frequency high imageability regular words, low frequency low imageability irregular words) with 25 items each. Imageability ratings were drawn from the Cortese & Fugett^14^ estimates of monosyllabic English words. Frequency was measured in SUBTLEX-US^15^ frequency per million words based on the Zipf distribution. As described in the Supplementary Material of Staples et al., 2025^9^, high and low frequency words significantly differed in mean frequency per million (*t*(99.03) = 6.57, *p* < .001) and were matched on mean letter length (*t*(187.34) = -0.28, *p* = 0.78), mean imageability rating (*t*(196.50) = 1.27, *p* = .20), mean articulatory complexity (*t*(187.86) = -0.71, *p* = .48), and proportion of irregular words. Regular and irregular words were matched on mean letter length (*t*(193.16) = 0.65, *p* = .51), mean frequency per million (*t*(189.02) = -0.40, *p* = .69), and mean imageability rating (*t*(198.00) = 0.03, *p* = .98), but irregular words had lower mean articulatory complexity (*t*(197.63) = -3.35, *p* < .001)^9^. High and low imageability words differed significantly in mean imageability rating *t*(181.27) = 25.17, *p* < .001), and were matched on mean letter length (*t*(196.80) = 0.28, *p* = .78), mean frequency per million (*t*(180.63) = -1.10, *p* = .27), mean articulatory complexity (*t*(191.99) = -0.31, *p* = .76), and proportion of irregular words^9^. Participants were instructed to read the word aloud as quickly and accurately as possible. Words were presented in two sets of 100 items with other non-reading tests administered between sets. Each set of 100 items was presented in four blocks of 25 items with optional breaks between blocks. Items were presented in a fixed order in order to perfectly balance the order of word types. Each block of 25 items started with a different word type, and the 24 remaining trials of each word type (those that did not start a new block) were preceded by items of each of the eight word types exactly three times. This fixed ordering was chosen to balance the sequence of item types to address prior evidence for sequential item effects in oral reading^16–18^.

### Pseudoword Oral Reading

Participants read aloud 80 monosyllabic pseudowords, all 3-6 letters in length. As described in Dyslin et al. (2025)^19^, 60 of these pseudowords represent three groups of pronounceable nonwords (20 stimuli each) that vary by whether corresponding orthography-to-phonology body mappings exist in English. 0M pseudowords (e.g., *“dofe”*) contain orthographic bodies that do not occur in the English lexicon^20^. 1M pseudowords (e.g., *“bink”*) have a single plausible pronunciation based on existing orthographic body correspondences (e.g., *-ink* as in *“sink”*). MM pseudowords (e.g., *“chead”*) permit two or more plausible pronunciations due to multiple existing body correspondences (e.g., *-ead* pronounced as in *“bead”* or *“head”*). Pseudowords were matched to words on body consistency, positional bigram frequency, and articulatory complexity^21^. Specifically, 1M pseudowords were matched to regular real words, and MM pseudowords were matched to irregular real words, in both the real word reading and lexical decision tasks. Pseudoword stimuli were adapted and extended from Friedman et al. (1992)^22^.

In the BUILD protocol, the remaining 20 pseudowords were short items matched to real words that differ by one initial consonant (e.g., *“tub”* versus *“mub”*). In the ReadMap protocol, the remaining 20 pseudowords were pseudohomophones. Pseudohomophone stimuli were matched to 20 of the real word stimuli described above on number of phonemes, *t*(35.49) = 0.12, *p* = .904, and the corresponding real words (e.g., *“box”* for *“bocks”, “hope”* for *“hoap”*, etc.) were matched to high-frequency real words on log word frequency, *t*(118) = 0.09, *p* = .929. Five participants completed a pilot version of the tasks in which the pseudohomophone “*plede*” was presented in place of “*bord*” for the pseudoword oral reading and lexical decision tasks. Data on the oral reading of the matched real words is also included for individuals who completed that test version. The matched pseudowords and real words are described further in Dickens et al. (2021)^8^.

Pseudoword stimuli were presented in blocks of 20 items, with breaks provided between each block. The order of stimuli presentation also varies by task version, with the BUILD version presenting items blocked by word type and the ReadMap version presenting words in a pseudorandomized order. Pseudowords were administered prior to the real word reading task in order to avoid word priming effects. Scoring was based on the first complete reading attempt that included both a consonant and a vowel. Responses were scored as correct if they contained a plausible pronunciation of the presented pseudoword. Pseudoword responses were coded as incorrect if they included spelling-to-sound mappings that do not occur in American English.

### Lexical Decision

Participants completed a classic visual lexical decision task. Two nearly identical versions of the task were administered, indicated by the “ProjectName” column. The BUILD version tested 400 items (200 words, 200 pseudowords), including the same 200 real words used for the oral reading task, the 60 pseudowords manipulated on spelling-sound mappings described above, 20 pseudohomophones, 60 pseudowords derived by altering regular words from the real word corpus, and 60 pseudowords derived by altering irregular words from the real word corpus. The ReadMap version tested 320 items (160 words, 160 pseudowords) in order to reduce testing time and improve matching across word types. The 160 words were a subset of the 200 oral reading items (20 of each word type), and the 160 pseudowords included the 60 items manipulated on spelling-sound mappings described above, the 20 pseudohomophones used for oral reading in the ReadMap protocol, 40 items derived by altering regular words from the real word corpus, and 40 items derived by altering irregular words from the real word corpus. Ten pseudohomophones tested in lexical decision overlapped between the BUILD and ReadMap task versions, and 10 differed. Overall, 310 items overlapped across the two task versions (160 words, 150 pseudowords). In this dataset, we have included trial-level data for all items tested, but to ensure comparability across all participants, we calculated summary statistics only for the shared items.

Each trial began with a 500-millisecond fixation cross, followed by stimulus presentation with response options displayed at the bottom of the screen. Participants indicated whether the stimulus was a “real word” or “not a word” by pressing the corresponding on-screen button. The task was administered in two blocks of 200 trials in the BUILD version or 160 trials in the ReadMap version. Response time (RT) was defined as the interval between stimulus onset and button press, and only RTs from correct trials are included in the data.

### Semantic Decision fMRI task

To identify language-sensitive cortex, participants performed an adaptive semantic decision fMRI task^11^. In the semantic condition, participants viewed pairs of words and indicated if they were related in meaning (e.g., calendar-date) with a button press. In the control condition, participants viewed pseudofont pairs (e.g., ∂DʊΣD–∂θʊƧD and indicated if they were identical. Ten alternating blocks of each condition were acquired. An adaptive staircase procedure adjusted the difficulty of the task based on performance independently for the semantic and perceptual blocks: when two successive correct responses are obtained in a condition, the task increased to the next difficulty level. When participants make an incorrect response, they move down two levels. Difficulty was modulated by manipulating lexical frequency, age of acquisition, word length, degree of relatedness, concreteness, and presentation rate for the semantic blocks and visual similarity of mismatching pairs for the perceptual blocks. To equate task demands, the presentation rate was adjusted at the start of each semantic block and held constant across the following pseudofont block.

### Neuroimaging Data Acquisition

All MRI data was acquired at Georgetown University’s 3T Siemens MAGNETOM Prisma scanner using a 20-channel head coil. Two structural images were acquired. First, a T1-weighted magnetization prepared rapid gradient echo (MPRAGE) sequence was acquired: 176 sagittal slices; slice thickness = 1 mm; 1 mm^3^ voxels; field of view (FOV) = 256 mm; matrix = 256 × 256 mm; flip angle = 9º; generalized autocalibrating partial parallel acquisition = 2; repetition time (TR) = 1900 ms; echo time (TE) = 2.98 ms; scan time: ∼ 5 mins. Second, a fluid-attenuated inversion recovery (FLAIR) sequence was acquired: 192 sagittal slices; slice thickness = 1 mm; 1 mm^3^ voxels; FOV = 256 mm; matrix = 256 × 256 mm; flip angle = 120º; generalized autocalibrating partial parallel acquisition = 2; TR = 5000 ms; TE = 386 ms; scan time: ∼ 6 mins. To assist with registering the BOLD data to the anatomical, a coplanar T2 was also acquired: 48 slices; TR = 4010 ms; TE = 76 ms; slice thickness = 2.6mm with a 10% gap; 2.9 mm voxels; FA 150°.

To assess language activation, 504 volumes of a BOLD T2*-weighted scan were acquired: 48 slices; TR = 794 ms; TE = 30 ms; slice thickness = 2.6 mm with a 10% gap; 2.9 mm voxels; FOV= 211 mm; matrix = 74 × 74; FA 50°; SMS = 4; scan time ∼ 7 mins. Spin echo field maps were acquired to correct magnetic field distortions: 48 slices; TR = 3130 ms; TE = 33 ms; slice thickness = 2.6 mm with 10% gap; FOV = 211; matrix = 74×74; FA = 90°; scan time: ∼7 mins. Scans were warped to the MNI152 template using Advanced Normalization Tools^21^.

To quantify white matter connections, multi-shell high angular resolution diffusion imaging (HARDI) scans were acquired: 74 axial slices; slice thickness = 2 mm; diffusion-weighted gradients: 81 directions at b = 3000, 40 at b = 1200, 7 at b = 0; 2 mm^3^ voxels; flip angle = 90º; phase encoding = anterior to posterior; partial Fourier = 6/8; FOV = 232 mm, matrix = 116 × 116 mm; TR = 4700 ms; TE = 82 ms; readout time = 61 ms; slice acceleration = 1; scan time: ∼ 10 mins). Six reverse phase-encoded b = 0 images were acquired for susceptibility field estimation (scan time: ∼ 1 min). To correct distortion artifacts, a blip up/down sequence was acquired with the same parameters as the HARDI sequence, except for acquiring both anterior-to-posterior and posterior-to-anterior phase encodings.

### Neuroimaging Data Pipeline

For all BOLD data, standard preprocessing was performed using AFNI^24^. Steps included realignment to account for motion, scaling to a global mean of 10,000, smoothing with a 6mm FWHM kernel, temporal high-pass filtering at 0.01 Hz, and detrending. Preprocessed BOLD images were then coregistered to the anatomical via the registered coplanar T2 image. A whole-brain general linear model was estimated using the *fmrilm* function from FMRISTAT^25^. The time-course of a white matter and CSF seed and the 6 head-motion parameters not convolved with the hemodynamic response function (HRF) were included as covariates. The task was modeled as two alternating boxcar functions, one for the Semantic task and one for the Pseudofont task, convolved with the HRF. A contrast of Semantic greater than Pseudofont task was estimated, and the resultant statistical maps were warped to the MNI152 template using the transformation computed from the MPRAGE.

HARDI data was processed using MRtrix 3.0^26^. Preprocessing included the standard stepwise application of Gaussian noise removal, Gibbs ringing artifact removal, correction of distortions induced by motion, eddy currents, and magnetic susceptibility, and inhomogeneity distortion correction. Voxelwise fiber orientation distributions were calculated from the HARDI data using multi-shell, multi-tissue constrained spherical deconvolution^27^. White matter pathways were estimated using probabilistic anatomically constrained tractography^28^ on the white matter fiber orientation distributions traced 15 million streamlines in native space (algorithm = iFOD2, step = 1, min/max length = 10/300, angle = 45, backtracking allowed, dynamic seeding, streamlines cropped at grey matter-white matter interface). The Spherical Deconvolution Informed Filtering of Tractograms-2 algorithm^29^ was applied to derive cross-sectional multipliers for each streamline to adjust streamline densities to be proportional to the underlying white matter fiber densities. White matter connectivity matrices were created for each participant by assigning streamlines to brain parcels of the Brainnetome atlas^30^. Each edge of the derived connectomes represents the apparent fiber density of the white matter connecting the two brain parcels.

## Data Record

The data set contains behavioral and neuroimaging data collected from neurotypical participants who completed the reading and language tasks as described in the Methods section above. All data is publicly available on the Open Science Framework (OSF; <u>link here</u>).

Behavioral data are organized into task-specific folders: Real Word Oral Reading, Pseudoword Oral Reading, Lexical Decision, and Semantic Decision. Each task folder contains behavioral data files and associated experimental scripts.

Behavioral data are provided in .csv format for trial-level data, and subject-level and item-level summary scores. Column headers for all behavioral files are defined in an accompanying “ReadMe” document. In addition, a document entitled “Appendix_Oral Reading Error Coding” provides a detailed description of the procedures used to classify oral reading responses. Task implementation files (E-Prime format) are included within each task folder for reproducibility purposes. Montreal Cognitive Assessment (MoCA)^12^ scores for each participant are also available in .csv format.

The included neuroimaging data consists of per-participant deidentified raw MRI scans, whole-brain connectomes derived from the HARDI data, and task-based activation maps derived from the Semantic Decision task. Neuroimaging files are organized at the participant level following standard conventions (Brain Imaging Data Structure (BIDS) specification^23^).

### Technical Validation

#### Behavioral task reliability

Split-half reliability for the behavioral reading tasks was estimated in R version 2025.5.0.496 using a resampling procedure. For each measure, items were randomly divided into two halves 1,000 times. Participant-level scores were computed for each half and correlated across halves, with reliability estimates corrected using the Spearman-Brown formula. In tables 2-4 below, we report the mean split-half reliability, standard deviation across iterations, and 95% resampling interval for each of the behavioral reading tasks. In the lexical decision task, the splits were stratified to separately balance real words and pseudowords. Overall, results indicate high reliability across measures for all tasks.

**Table 2.**
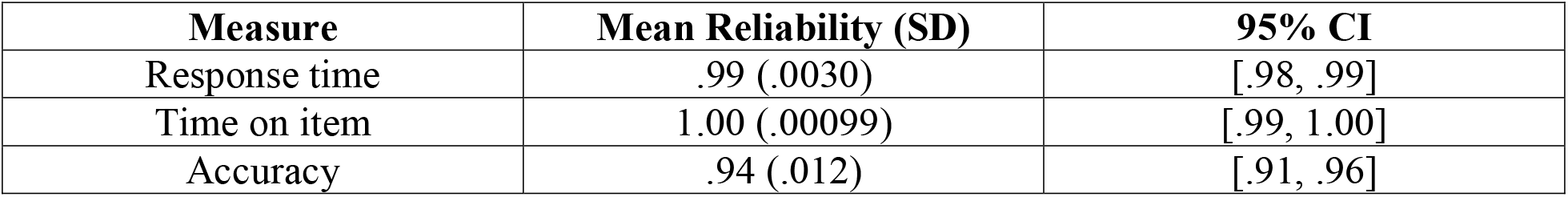
Real word oral reading split-half reliability estimates.

**Table 3.** Pseudoword oral reading split-half reliability estimates.

| Measure | Mean Reliability (SD) | 95% CI |
| --- | --- | --- |
| Response time | .99 (.0058) | [.97, .99] |
| Time on item | .99 (.0028) | [.99, 1.00] |
| Accuracy | .97 (.0071) | [.95, .98] |

**Table 4.**
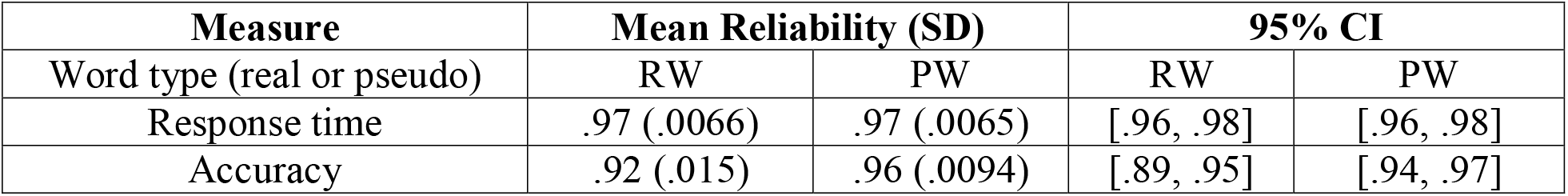
Lexical decision split-half reliability estimates. Estimates were calculated separately for real words (RW) and pseudowords (PW).

#### fMRI Quality Control

We evaluated head motion and the temporal signal-to-noise (tSNR) ratio to assess the quality of the minimally preprocessed fMRI data (Fig. 2). Head motion was quantified using framewise displacement (FD)^24,25^. All participants save two average less than 0.5mm mean FD, a commonly used cutoff ^24^, with a third participant just below. We computed tSNR as the voxelwise mean activation divided by the standard deviation of the voxel time series, after motion correction^26^. For visualization purposes, tSNR maps were warped to the MNI152 template. The data showed good tSNR in language-critical frontal, temporal, and parietal regions.

**Figure 1.**
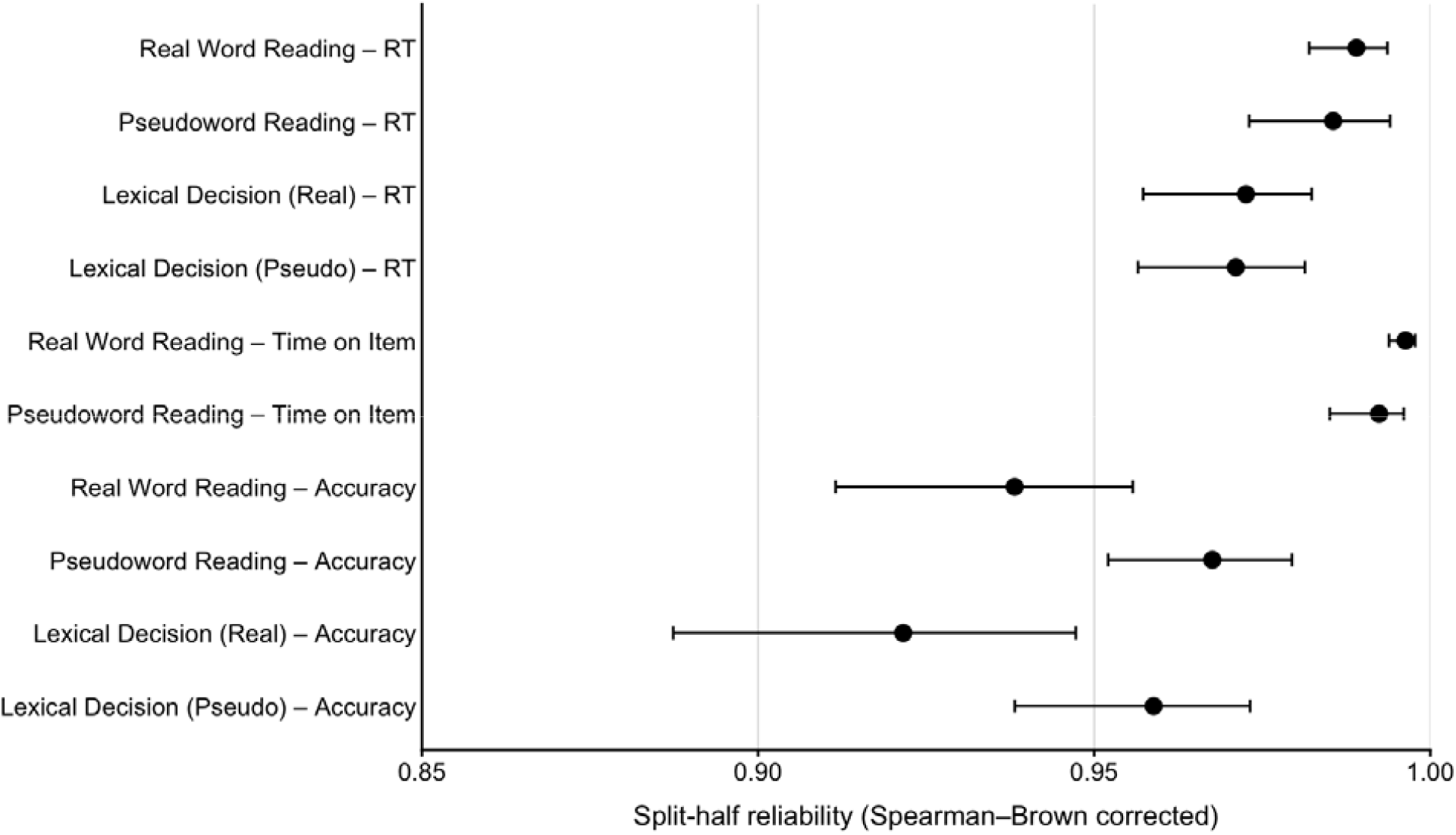
Dot and whisker plot depicting the mean estimate and 95% resampling interval for each measure per task. Real words and pseudowords were sampled separately for the lexical decision task.

**Figure 2.**
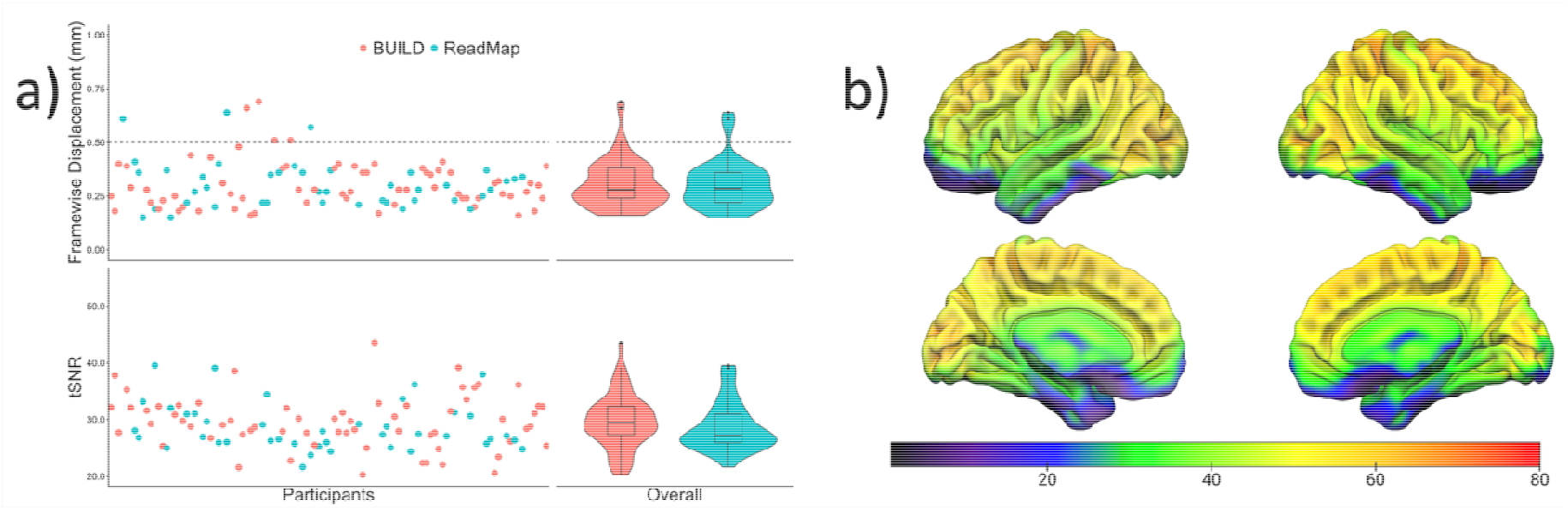
Minimally preprocessed fMRI quality control metrics. a) The participant-wise and mean framewise displacement and temporal signal-to-noise (tSNR) values for every participant in the sample, color-coded by project. b) The mean tSNR values across the sample, projected onto the MNI152 template. Overall, the data show low motion and good sensitivity to signal.

To evaluate the quality of the processed data, we computed the mean of the Semantic > Pseudofont contrast across all participants and compared it to an automated “association” meta-analysis for the term “language”, created using NeuroSynth^27^ (https://neurosynth.org/). NeuroSynth automatically extracts activation coordinates and keywords from published neuroimaging papers. The association test then compares whether activation in a region is more associated with studies tagged with a given keyword (here, “language”) than those that are not tagged with that keyword (Fig. 3). Overall, activations from the semantic decision task overlapped strongly with the core language regions identified in the NeuroSynth meta-analysis, alongside regions associated with semantic processing such as the anterior temporal lobe and the angular gyrus.

**Figure 3.**
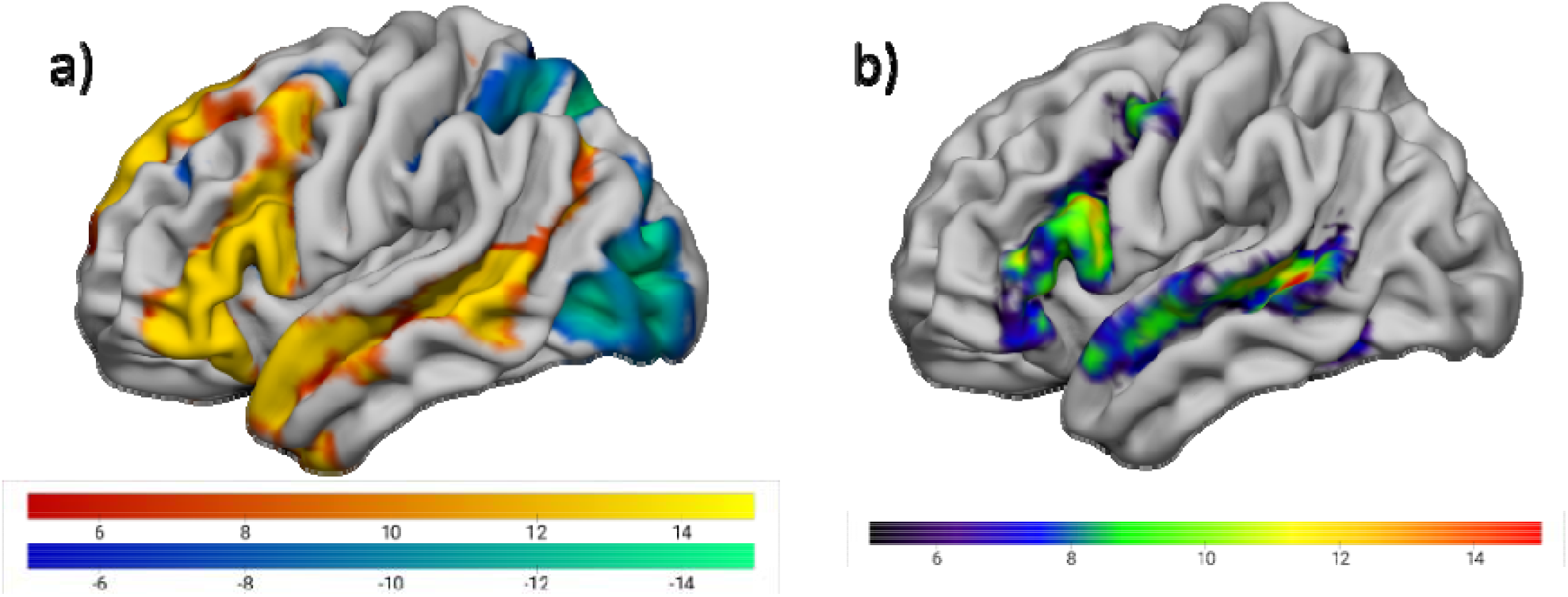
Processed fMRI quality control projected onto the MNI152 template. a) The average Semantic > Pseudofont contrast across the sample. Hot colors indicate areas more activated by the Semantic condition than the Pseudofont condition. b) The result of an automated meta-analysis for “language” derived via NeuroSynth. The semantic decision task identified core areas associated with language processing.

**Fig. 4.**
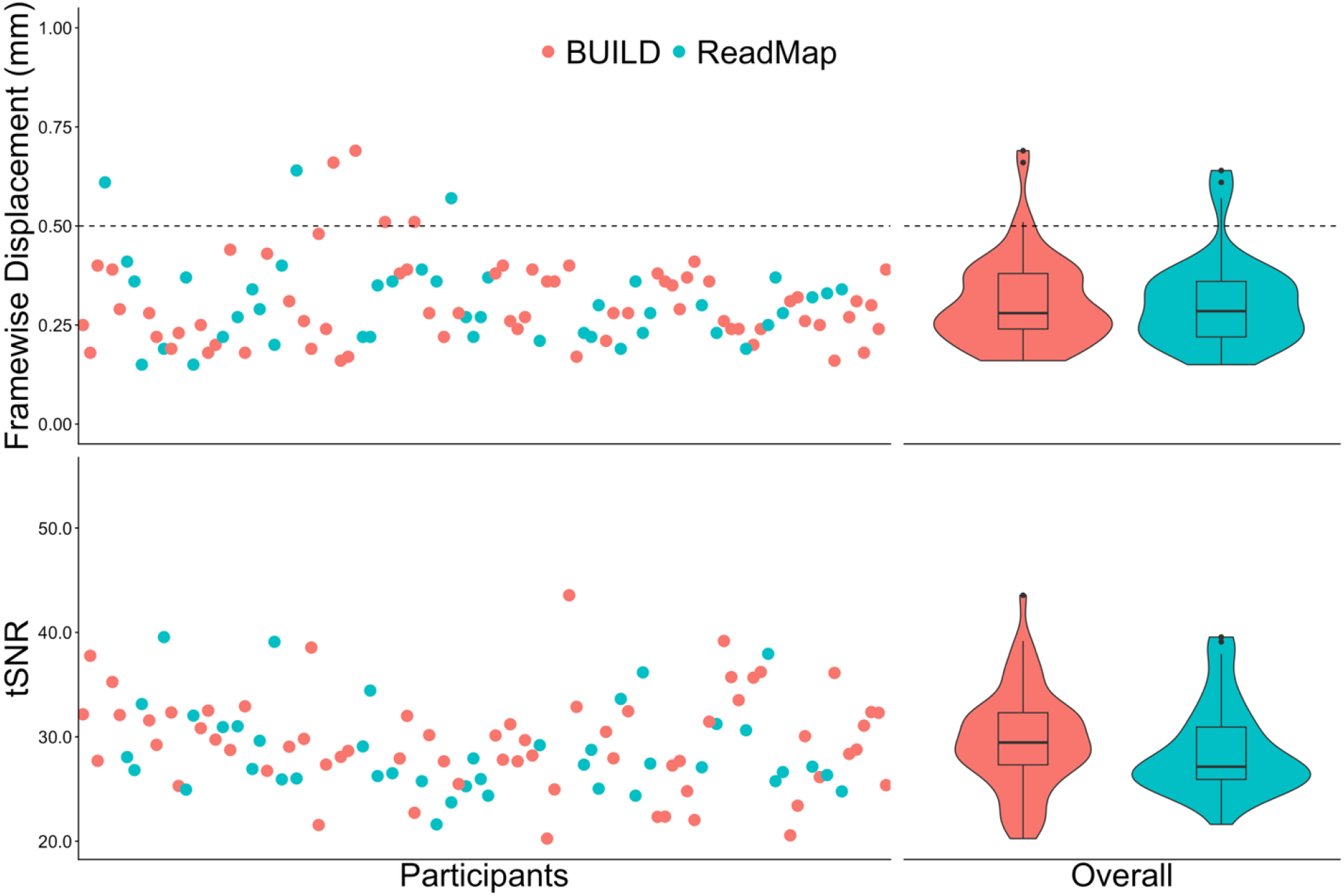
DTI quality control metrics. Overall, the DTI data shows low motion and good signal sensitivity.

#### HARDI Quality Control

The quality of the HARDI data was assessed using FSL Eddy^28,29^. We assessed the same metrics as in the task-based fMRI data: FD and tSNR (here, at *b* = 0). Only seven participants were above the commonly used cutoff of 0.5mm mean FD^24^.

## Data Availability

All data described has been deidentified and made publicly available via the Open Science Framework (OSF; <u>link here</u>). The repository includes behavioral data (trial-, item-, and subject-level), E-Prime scripts, and neuroimaging data (structural scans, connectomes, and task-based activation maps). No restrictions apply to data access. All files can be downloaded directly from the OSF repository.

## Code Availability

E-prime files and neuroimaging preprocessing scripts are available via OSF (<u>link here</u>).

## Acknowledgements

We wish to thank the individuals who contributed to data collection, in alphabetical order: Davetrina Seles Gadson, Catherine “Trini” Kelly, Elizabeth Lacey, Sarah Snider, and Candace van der Stelt.

## Author Contributions

Experiment design: P.E.T., J.V.D., S.M.D.

Data Collection: J.V.D., D.M., S.P., E.D.

Original Draft Preparation: E.A., R.S., S.M.D., E.H.T.C., A.L.

Review and Editing: E.A., A.L., P.E.T.

